# Freebie: A Multi-Camera System for Real-Time Markerless Hand Tracking in Non-Human Primates

**DOI:** 10.64898/2026.11.11.765959

**Authors:** J. Joseph, M. Ceradini, L. Liu, M. Kelberman, M. Mender, J. DeMatteo, O. Lee, H. Temmar, M. G. Copenhaver, A. Ward, T. Kung, C. Chestek, A. Draelos

## Abstract

**Objective:** Neural decoders for dexterous finger movements are limited by the accuracy with which ground truth movement is measured. In non-human primates (NHPs), hand kinematics are typically recorded with instrumentation that constrains movement dimensionality. We evaluated whether a markerless camera system could replace conventional methods for tracking movements in real time.

**Approach:** We developed Freebie, a real-time markerless system for tracking NHP finger movements. Keypoints detected online were used in place of manipulandum bend sensor signals. Two NHPs performed tasks by flexing fingers to move a virtual hand into target zones. The virtual hand was directly controlled using bend sensors or Freebie-derived kinematics, or indirectly controlled through a Kalman filter (KF) decoding kinematics from forearm electromyographic (EMG) activity. We quantified Freebie’s accuracy relative to manual labels, agreement with bend sensor measurements, and latencies. We assessed online performance under Freebie control and compared EMG KFs trained using Freebie-derived labels with those trained using bend-sensor-derived labels.

**Main Results:** We observed a median end-to-end latency of 15.8 ms at a 30 FPS camera acquisition rate, and 16.2 ms at 60 FPS. Freebie labels closely tracked bend sensor positions (median |r|=0.96). Median keypoint errors were 1.7 to 6.5 pixels, below the 20-pixel width of the smallest tracked finger. Direct Freebie control did not differ from bend sensor control on five of six online metrics; path efficiency was higher with Freebie. EMG KFs trained on Freebie-derived labels performed similarly to those trained on bend sensor labels online. When evaluated offline, models trained with Freebie had lower error.

**Significance:** These findings indicate that bend sensors are not required to obtain finger labels for training a decoder: markerless keypoints alone provided effective labels. This is an important first step towards real-time tracking of more complex movements, beyond what could be easily measured with a manipulandum.

## 1. Introduction

Over the last decade, intracortical brain-machine interfaces (BMIs) have progressed from basic cursor control to real-time control of individual finger movements [1]. In non-human primates, (NHPs), linear and neural network decoders can predict simultaneous, independent movements of two finger groups from intracortical spiking activity [2,3] and these signals have been integrated with closed-loop functional electrical stimulation (FES) to restore continuous finger function in temporarily paralyzed hands [4,5]. In human clinical trials, intracortical recordings have further revealed compositional representations of complex, multi-joint movements [6,7], enabling continuous real time control across up to four degrees of freedom (DOFs) in three independent finger groups [8]. Together, these findings establish that intracortical signals supporting high-dimensional, continuous hand control can be recorded and decoded.

The quality and dimensionality of behavioral labels are critical to training and evaluating such decoders. A decoder’s ability to learn a transformation from neural activity to complex movement is in part limited by the accuracy with which the movement itself is measured. In nearly all NHP hand studies, kinematics are captured with constrained, low degree of freedom (DOF) instruments. Common examples include center-out joysticks for gross movements, touchscreens, and, in our group, manipulanda equipped with bend sensors to track one or two finger-groups while the hand is held in a fixed position [2,9]. Marker-based optical motion capture, data gloves, and instrumented goniometers can capture additional dimensions but require physical attachment, extensive training, repeated recalibration, and most importantly, they restrict natural movement in NHPs. Behavioral measurement, rather than neural recording, often limits the complexity of movements that can be studied and decoded [3,10]. These constrained tasks also narrow the range and diversity of observed hand movements, thus limiting insights into naturalistic motor control [11].

Markerless pose estimation offers a direct method for overcoming these limitations. This approach can recover high-DOF hand and body kinematics without physical markers or constrained postures, approaching the classical 27-DOF articulated hand model [12] used in state-of-the-art human hand-tracking systems [13]. Robust deep neural networks pretrained on large datasets can track user-defined body parts in conventional video with accuracy comparable to that of human annotators after training on only a few hundred labeled frames [14–16]. Extensions support three-dimensional (3D) reconstruction from multiple cameras [16–18] including systems developed specifically for macaques [19–21]. Post-processing methods, including Kalman smoothing [16,17] and biomechanical constraints [22], can further improve the reliability of tracking occluded or uncertain keypoints.

However, most markerless tracking tools were built for offline analysis of recorded videos and were not designed to deliver kinematic estimates inside a real-time control loop nor for NHP hands. Their real-time operation is limited by the computational and memory-intensive demands of multiple camera views, long inference times of large neural networks, and non-causal post-processing methods. Notable exceptions include DeepLabCut-Live! (DLC), which reports pose estimation latencies below 15 ms and processing rates exceeding 100 frames per second (FPS) [23], and SLEAP, which reports inference latencies below 3.5 ms [15]. Nevertheless, these systems typically operate on single-camera views, which do not allow for 3D reconstruction, and have primarily been demonstrated for triggering simple feedback events. To our knowledge, no published system performs synchronized multi-camera markerless inference with optional 3D kinematic reconstruction while bidirectionally exchanging kinematic and additional experimental data with external hardware (including neural signal processors, real-time control computers, and stimulators) within the tens of milliseconds latency required for closed-loop BMI applications.

We sought to close this gap by integrating existing validated components into Freebie, a novel real-time, multi-camera, markerless hand-tracking system for NHPs. Our system currently integrates up to 4 synchronized industrial cameras, a real-time PyTorch model built from a DLC pose estimation base, and the *improv* real-time software framework [24] with a Redis-backed message queue to coordinate image acquisition, inference, visualization, and hardware input and output (I/O). We evaluated model accuracy relative to human annotations and compared markerless estimates with measurements from existing bend sensors. To test whether markerless keypoints can serve as training labels, we trained a Kalman filter predicting the tracked finger movements from EMG recorded from implanted electrodes in the forearm. The decoder architecture and finger task were similar to those used in previous intracortical BMI studies [2,9]. The source of the training labels was the only differing factor between conditions, isolating the effect of label quality from the choice of input signal. Finally, we quantified system latency and throughput to evaluate Freebie’s use in real-time applications, including online closed-loop hand control and training of an EMG decoder. We demonstrated extensions to 3D tracking during free behavior and FES characterization experiments.

## 2. Methods

### 2.1 Subjects, implant, and ethics

The University of Michigan Institutional Animal Care and Use Committee approved all procedures (PRO00011815), which were performed in accordance with the NIH Guide for the Care and Use of Laboratory Animals. Two adult male rhesus macaques (*Macaca mulatta*), Monkey R (18 years and 5 months) and Monkey N (13 years and 5-9 months), participated. Forearm electromyography (EMG) was recorded from both NHPs using implanted chronic bipolar intramuscular leads (Synapse Biomedical, Oberlin, OH, USA). Wires were surgically implanted into the forearm muscles and tunneled to an interscapular exit site where they were housed in a protective primate jacket (Lomir Biomedical, Inc., Malone, NY, USA). Respectively, Monkeys N and R were implanted with 8 and 16 electrodes (Fig 1A), respectively, into extrinsic flexor and extensor muscles that control finger and wrist function. While Monkey R was implanted with 16 bipolar electrodes, only eight bipolar EMG channels used for decoding were selected, as Monkey R’s flexor electrodes were unexpectedly disconnected from their leads, without causing any harm. Each electrode was implanted by making an intramuscular tunnel within the muscle belly. The electrode was then buried into the muscle tissue, ensuring that all contacts were completely enveloped and secured with monofilament sutures. EMG data was analyzed 1790-1903 days and 1199-1210 days after intramuscular implantation. Results in the main text are from Monkey R, recorded with the current version of the system. Monkey N was recorded with an earlier version of the system (Section 2.4), which is reported in the Supplementary Material. We additionally demonstrate Freebie multi-DOF hand control using Freebie 3D tracking in one human (Supplementary Fig. S1).

**Figure 1.**
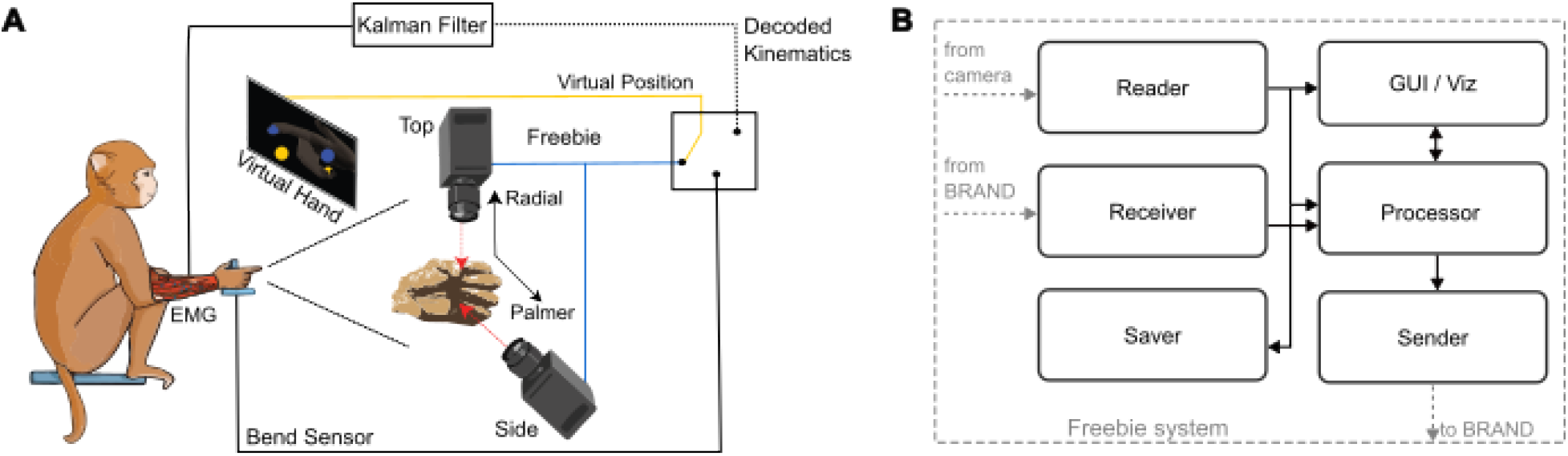
Hardware and software architecture. (**A**) Closed-loop flow of information during an experimental task. The NHP moved his hand in a manipulandum while cameras acquired images of the hand and fingers in real time. Tracked finger positions (blue) or bend sensor values (solid black) controlled a virtual hand (yellow) displayed on a monitor. Concurrently, EMG activity recorded from intramuscular flexor electrodes implanted in the left forearm were decoded into predicted kinematics (dashed black) that serve as an alternative control signal for the same virtual hand. **(B)** Graph of data flow among actors in the processing pipeline. Images were acquired by a camera reader and passed to the processor, video saver, and graphical interface nodes; processor outputs containing keypoint estimates were passed to the sender.

### 2.2 Signal Processing

For both NHP subjects, percutaneous EMG leads were connected to a Cerebus neural signal processor (NSP, Blackrock Neurotech, Salt Lake City, UT, USA) via a custom connector manufactured by Synapse Biomedical (Oberlin, OH, USA). The electrodes were connected to the data acquisition system through a splitter box receiving TouchProof connections and a grounded reference electrode. The reference electrode was attached to an implanted head-post by an alligator clip [1]. Incoming Cerebus data was sampled at 30 kHz and a digital offset removal high-pass filter (0.25 Hz) was applied by the pre-amplifier, and the signals were band-pass filtered between 100 and 500 Hz using a 4^th^-order Butterworth filter before being downsampled to 2 kHz and summed into 32 ms bins. For Monkey R, these signals were then streamed over UDP to a BRAND real-time computer [21] which ran the task, the bend sensor and Freebie inputs, and the decoder. For Monkey N and the earlier Freebie system, signals were instead sent to a different real-time computer (xPC Target, Mathworks, Natick, MA, USA) which operated at a 1-ms update interval.

### 2.3 Experimental rig and behavioral task

During all sessions, the monkeys were head-fixed and their arms were comfortably restrained at a resting right angle. The monkey’s left hand was placed in a custom manipulandum instrumented with bend sensors that measured finger and wrist flexion (FS-L-0073-103-ST, Spectra Symbol, Salt Lake City, UT), as described previously [2,9]. Bend sensors were sampled at 1 kHz and calibrated daily in a normalized range from 0–1. A monitor provided visual feedback by displaying a virtual hand and targets (MusculoSkeletal Modeling Software [26]) (Fig. 1A).

Monkey R performed a one-DOF index finger task on a BRAND operated rig. Bend sensor voltages were digitized by an Arduino UNO R4 (14-bit ADC) and streamed at 460,800 baud. The newest sample was published at 200 Hz, normalized from 0 (extended) to 1 (flexed) using a per-run calibration range, clipped, and smoothed with a 20-sample (100 ms) moving average. Each trial presented one target drawn uniformly from 0–1 on the index DOF, and all four fingers of the virtual hand followed the index DOF. Monkey N performed one-DOF and two-DOF (index and MRS moving separately) tasks on an xPC Target system, based on previous tasks described in [2,6]. A trial was successful when the virtual hand stayed within the target radius (0.10 on the first recording day and 0.15–0.20 thereafter) for the hold time (0.5 or 0.75 s), and timed out after 10 s, with no inter-trial interval. Each success was rewarded with 150 ms of juice, and the task was scored at 100 Hz.

The virtual hand was driven in one of three ways: (i) bend sensor control, in which the bend sensor signal directly controlled the virtual hand; (ii) Freebie hand control, in which the bend sensors were disconnected and the Freebie estimate of finger positions directly controlled the virtual hand; or (iii) closed-loop EMG decoding, in which a Kalman filter trained on either Freebie-derived or bend sensor-derived labels controlled the virtual hand based on EMG activity in real time (Fig. 1A) (Section 2.10). For Monkey R, the control source was selected in software by a digital switch, and the bend sensor continued to be recorded during Freebie control. For Monkey N, the bend sensors were disconnected during Freebie control, and Freebie estimates were sent to the xPC over a serial connection.

### 2.4 Freebie real-time tracking system

#### Hardware and framework

Freebie used the *improv* framework as a backend system [24]. With *improv,* separate processes called “actors” were used to manage frame acquisition, pose inference, serial output to the task computer, packet reception, video saving, and display. Actors exchanged data through a shared in-memory store (Fig. 1B). Data from each camera was acquired through separate reader actors acting in parallel. All cameras were industrial color cameras (The Imaging Source DFK 33UX252). For Monkey R, a single camera viewing the index finger from the front left delivered 960 × 720 RGB frames at 30 FPS on one recording day and 60 FPS on the remaining days, with exposure fixed at 2 ms, gain 0, and fixed white balance. For Monkey N, a top camera viewing the index finger and a side camera viewing the MRS finger group recorded 1920 × 1080 RGB frames at 30 FPS, with automatic exposure, gain, and white balance. The cameras were connected via a fiber-optic link to a Linux workstation (Intel Core i9-14900K, 32 threads; 128 GB RAM; NVIDIA GeForce RTX 4090, driver 550.163; Ubuntu 24.04 LTS). In this study, Freebie was evaluated using configurations of one, two, and four cameras. Multiple light sources were placed around the workspace to minimize shadows on the fingers. Frames reached each camera actor through a GStreamer appsink with a five-frame queue. When the queue was full, the oldest frame was dropped, so the newest frame was always the one processed. Each frame was timestamped using the acquisition computer’s clock, JPEG-encoded (Monkey N configuration), and written to the shared store. The frame’s store key, timestamp, and frame number were then passed to the processor, sender, and video-saver actors.

#### Keypoint detection

Keypoints were estimated using a model based on DLC (DeepLabCut 3.0, PyTorch engine) [14], with ResNet-50 backbones, group normalization, and a heatmap head with location refinement. The inference class from DLC was modified for Freebie to run online in a lightweight PyTorch actor that avoided re-instantiation with each image.

For Monkey R, a single model tracked one keypoint, the index PIP joint, in the front-left camera view. It was trained on 80 manually labeled frames (20 from each of four videos recorded on three days before the experiments) with a 95%/5% train/test split. Training for 200 epochs used AdamW (learning rate 5 × 10⁻⁴, reduced to 1 × 10⁻⁴ at epoch 90 and 1 × 10⁻⁵ at epoch 120), a batch size of 8, affine augmentation (probability 0.5, rotation up to ±30°, scaling 0.5–1.25). Data augmentation also included hybrid 448 × 448 crop sampling with a maximum shift of 0.1, Gaussian noise (σ = 12.75), and motion blur. The checkpoint with the lowest test error (epoch 10) was deployed, and the same model was used on all recording days.

For Monkey N, a separate model was trained for each of the two camera views. The top-view model tracked four keypoints: the distal interphalangeal joint (DIP), the proximal interphalangeal joint (PIP), the metacarpophalangeal joint (MCP), and the wrist. The side-view model tracked one MRS keypoint located at the middle fingertip. Black Sharpie marks were applied to the MCP joint and the wrist to increase visual contrast. The approach remained markerless because it used no physical or reflective 3D markers or instrumented sensors. Because the DIP and PIP joints were occluded by the manipulandum, their labels represented the inferred rather than directly visible joint locations (Fig. 2A). Monkey N’s models were trained on 514 (top) and 308 (side) labeled frames using the same training configuration above. The best-performing checkpoints were deployed: epoch 40 for the top view and epoch 10 for the side view. Human annotators labeled all frames, including a subset consisting of corrections to predictions from an earlier model rather than labels made from scratch.

**Figure 2.**
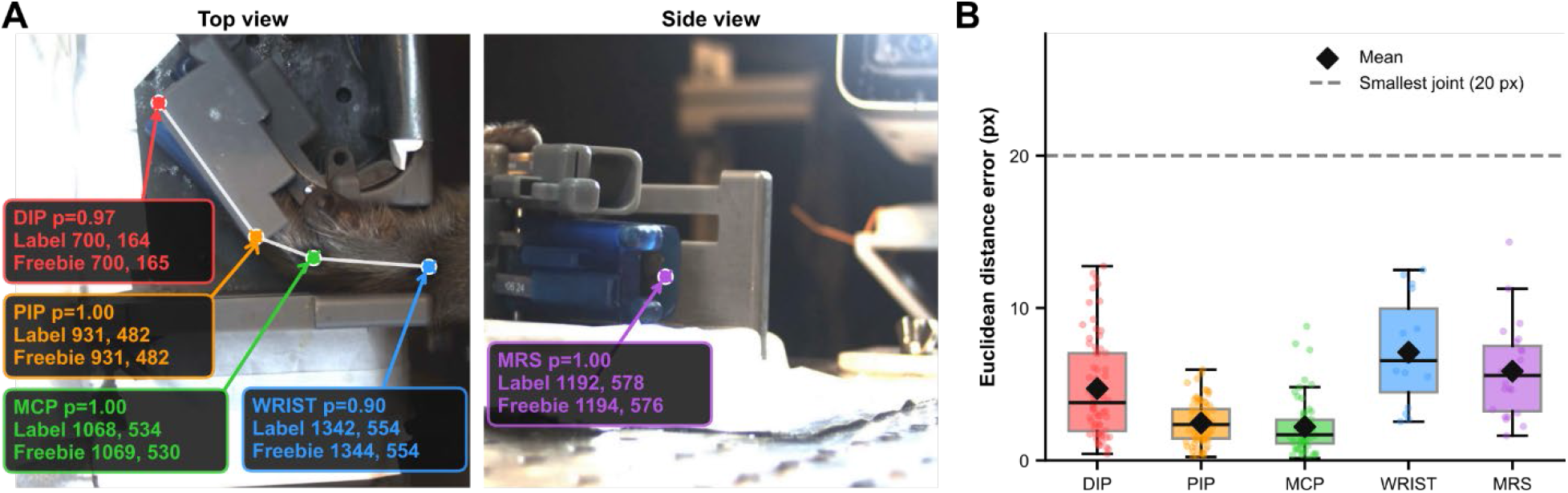
Freebie tracked joints with high accuracy. (**A**) Example held-out frames from top and side view cameras. Circles are manual annotations and crosses are Freebie predictions for the DIP, PIP, MCP, wrist and MRS keypoints. Labels give the annotated and predicted pixel coordinates and the prediction likelihood (p). (**B**) Euclidean error between manual annotations and Freebie predictions for each keypoint for held-out frames with prediction likelihood ≥ 0.99 (DIP n = 68, PIP n = 80, MCP n = 59, wrist n = 15, MRS n = 23 frames). Boxes show the median and interquartile range, whiskers extend to 1.5× the interquartile range, diamonds mark the mean, and dots are individual frames. The dashed line marks 20 px, the approximate diameter of the smallest labeled joint.

Monkey N’s online pipeline was developed over the course of the study, so earlier sessions used earlier versions. Modifications included the addition of an adaptive forward prediction to keypoints using Kalman filters, queue draining by discarding old frames, and replacing the index angle with the PIP y coordinate as the tracked keypoint of interest. After Monkey N’s last session, we optimized system latency by removing JPEG compression before images were written to the shared store, moving downsampling into the camera’s acquisition step, and assigning each acquisition and inference actor to a dedicated physical performance core of the host processor. Frames from multiple cameras were also batched into a single tensor and processed in one model forward pass. Monkey R’s experiments used this optimized pipeline with a single camera.

#### Video Saving

A video saver actor for each camera collected frame keys from the store into 5-s segments (150 frames at 30 FPS) (Fig 1B). A background thread wrote each segment to disk as a binary file of length-prefixed frames. Writing to disk was kept out of the real-time path. After each session, on request from the experimenter via the GUI, the segments were converted to one H.264 MP4 per camera with ffmpeg. Each segment was deleted once it had been successfully converted. The saver also logged the time at which it received each frame.

#### Graphical Interface

A PyQt5 display actor showed the live frame from every camera in a grid, redrawn at the camera frame rate (Fig 1B). The predicted keypoints and skeleton were drawn over each frame, along with the current joint angle. A pyqtgraph panel plotted the last 100 angle values for each camera.

#### Logging

Every actor logged wall-clock timestamps from the tracking computer for each respective process: frame-acquisition start, processor start and per-step durations, and the time of each output transmission. For Monkey N, keypoint predictions and all incoming xPC packets were also saved by the receiver actor (Fig 1B). Each packet carried bend sensor values and the xPC experiment time (eTime, ms) and was stamped with its arrival time on the tracking computer at ≈1 kHz. The xPC/Simulink model operated on a fixed 1 ms cycle. For Monkey R, the sender also logged each frame’s end-to-end latency, and BRAND logged all sensor, task, decoder and EMG streams (Section 2.5).

#### 3D Reconstruction

During free behavior experiments, we tracked 21 keypoints according to the Mediapipe landmark guide [12]. Our system is agnostic to the type of model used for keypoint extraction and is compatible for real time use with JARVIS-Mocap [27] and Mediapipe [13]. We calibrated using an 8 × 6 checkerboard with 25 mm squares and in later sessions with a ChArUco board (4×6, 42 mm squares, 31.5 mm markers, 4 marker bits, dictionary size 50) that was rotated and translated throughout the workspace while being recorded simultaneously by all cameras in the configuration. For 3D reconstruction, we estimated six extrinsic parameters per camera describing its position and orientation relative to the other cameras (three rotation and three translation). The internal coordinate system of each camera is described using intrinsic parameters (focal length, principal point and lens distortion). These were estimated using Anipose [17] and subsequently used for lens-distortion correction and 2D-to-3D projection. Because the cameras were rigidly fixed, calibration can be performed once and the resultant parameters reused across sessions. 3D keypoint positions can be obtained by triangulating the camera-specific 2D detections using a direct linear transformation (DLT).

#### Human 3D hand tracking

To test multi-camera, multi-DOF operation, one author familiar with the task performed the same random-target task with up to four controlled DOFs using real-time 3D hand tracking. Four cameras (960 × 720 RGB, 60 FPS, exposure 2 ms), calibrated with Anipose [17], were jointly processed by a single processor actor. At each step, the actor gathered the newest frame from each camera, waiting up to 12 ms, and used MediaPipe Hand Landmarker [13] on all images in parallel on the CPU to obtain 21 keypoints per hand. Detections were paired across cameras geometrically by triangulation with a direct linear transform (mean reprojection error < 8 px and wrist-to-middle MCP length of 20–200 mm), tracked across frames, and smoothed with a 3D Kalman filter. Because the cameras were not hardware-synchronized, the oldest frame contributing to a pose was a further 8–12 ms older than the newest frame, from which latency was measured. The 3D keypoints were streamed over UDP to BRAND, where a 27-DOF kinematic hand model [12] computed metacarpophalangeal (MCP) flexion angles. These were normalized to 0–1 using ranges from a guided calibration at the start of each run and mapped onto the virtual hand: index (one DOF); index and MRS (two DOFs); index, middle, ring– small (RS) (three DOFs); and index, middle, ring, small (four DOFs) (Fig. S1A).

### 2.5 Data processing

#### Sessions

Monkey R’s dataset comprised 32 runs recorded over 4 days: 15 bend sensor hand-control runs, 9 Freebie hand-control runs, and 8 closed-loop decoder runs recorded over 2 days (4 with a Kalman filter trained on bend sensor labels and 4 with one trained on Freebie labels). Monkey N’s dataset comprised 87 runs recorded over 24 days between 6 November 2025 and 27 February 2026, corresponding to 77 xPC runs. Each run’s data was converted to the Neurodata Without Borders (NWB) format.

#### Clock alignment

For Monkey R, all BRAND signals (bend sensor, Freebie, virtual hand, decoder output, EMG, trial events and control-source switches) were logged to a Redis database on a common clock, so no alignment was needed. Each run’s task window spanned the first trial start to the last trial result, and Freebie latency logs from the tracking computer were assigned to a run when their Unix timestamps fell within this window. For Monkey N, Freebie timestamps were mapped onto xPC time using an offset. The clock offset of a session was the median over all packets of (arrival time − eTime); the median was used because arrival stamps jitter around 1 ms per packet. The xPC run window on the tracking clock was defined as [first trial start, last trial stop] plus this offset. An offset and run window could be computed for the 41 sessions with both Freebie processor timestamps and xPC packets.

#### Offline re-inference

For some of Monkey N’s sessions, the saved videos were also re-analyzed offline with the final deployed model, because the live model used that day was underfit. The offline predictions replaced the live ones, per camera. This applied to 10 sessions for the top camera and 5 for the side camera. All other Monkey N sessions and all Monkey R runs used the live predictions.

#### Exclusion criteria

Before analysis, every session was audited for the data available to each analysis: NWB, trials, EMG, xPC packets, keypoints, frame timestamps and latency logs. Online metrics used closed-loop trials only. The bend sensor and Freebie agreement analysis required bend sensors connected to the xPC, received xPC packets and a known run window. Freebie-trained offline KFs required EMG, bend sensors and at least 20 s of joint coverage of all required Freebie features within the run. Two-DOF sessions without side-camera timestamps were analyzed on the index DOF only. Latency analyses used sessions with per-frame end-to-end logs, and the per-actor summary required at least 100 logged frames. The first logged frame of every session was discarded as GPU code initialization cost. After exclusion, the online-metric analyses included 51 bend-sensor and 17 Freebie hand-control runs (46 and 12 of them two-DOF, respectively), and 4 bend-KF and 4 Freebie-KF closed-loop runs (1 and 2 of them two-DOF, respectively).

### 2.6 Keypoint tracking accuracy

Tracking accuracy was calculated for Monkey N’s two-camera models on manually labeled frames: 597 top-view frames from 18 videos and 395 side-view frames from 13 videos. Each labeled frame was paired with the prediction from the same image. For each keypoint, frames with a prediction likelihood below 0.99 were discarded. The error was the Euclidean distance in pixels between the predicted and labeled positions. We compared this with a 20 px benchmark, as this was the approximate diameter of the smallest labeled finger width. We reported error on a held-out subset of 83 top-view and 87 side-view frames not used for training from 16 and 8 videos, respectively. This left 68 (DIP), 80 (PIP), 59 (MCP), 15 (wrist) and 23 (MRS) frames. For Monkey R’s single-keypoint model, two training shuffles of the same labeled data had test set RMSE of 2.95 and 2.77 px.

### 2.7 System latency

For Monkey R, end-to-end latency was measured for every frame as the time from the camera callback in the camera actor to transmission of the corresponding keypoint estimate over UDP, both on the tracking computer’s clock. Three stage times were also logged per frame: frame acquisition (the camera callback to the frame being queued for the processor), inference (the processor step, including the store read, DLC inference, the keypoint Kalman filter and post-processing) and sender (the sender step, on steps that sent a fresh prediction). The first frame of each session was discarded as GPU warm-up, and frames were kept when their timestamps fell within a run’s task window. End-to-end latencies were pooled across all 32 runs (690,485 frames) and compared with the 33.3 ms frame period of each run’s frame rate (33.3 ms at 30 FPS, 14 runs; 16.7 ms at 60 FPS, 18 runs). Stage times were averaged per run and summarized across runs. Monkey N’s latency was computed comparably from the earlier pipeline’s logs (20 sessions, 144,888 frames; Supplementary Fig. S2).

### 2.8 Freebie and bend sensor agreement

For each of Monkey R’s hand-control runs (n = 24), the Freebie signal (normalized PIP y coordinate after smoothing) was compared with the normalized and smoothed bend sensor signal. The bend sensor was linearly interpolated at each Freebie sample time at zero lag within the run’s task window, and the absolute Pearson correlation |r| was computed per run.

For Monkey N (Supplementary Fig. S3), for each bend sensor session with a known run window, each Freebie feature was compared with its corresponding bend sensor DOF: the index bend sensor for the index angle and PIP y coordinate, and the MRS bend sensor for MRS x coordinate. The bend sensor from the xPC stream (≈1 kHz) was linearly interpolated at each Freebie frame time. Only frames inside both the xPC coverage and the xPC run window were used. The Pearson correlation between the Freebie signal and the interpolated bend sensor was computed per session. Absolute values |r| were analyzed. Correlations were available for 30 sessions for the index finger signals and 16 for MRS x.

For Monkey N’s example traces (Fig. S3A), Freebie traces were mapped onto xPC time with the session offset and scaled robustly to 0–1 with the 20th and 80th percentiles of the total range of recorded values within the run and overlaid on the 1 kHz bend sensor for 8 consecutive trials. These trials started one third of the way through the successful, non-blank trials that lay entirely within the Freebie coverage.

### 2.9 Online performance metrics

The virtual hand position was calculated based on the bend-sensor or Freebie input (hand control) or based on the EMG decoder output. The success rate was the fraction of trials where the monkey placed his fingers within the target(s) for the hold time. Targets per second were defined as the number of all trials in the run divided by their summed duration. Time to target was the time from trial start to the first sample at which the distance between the virtual finger and the target center was smaller than the target radius. Orbiting time was defined as *t*_orbit_ = max (*N* Δ*t* −*t*_hold_ − *t*_target_, 0) where N is the number of samples in the trial, Δt is the sample period, t_target is the time to target, and t_hold is the hold time. Path efficiency was the straight-line distance between the first and last virtual hand positions divided by the summed sample-to-sample path length, with higher values corresponding to more direct straight routes. Bit rate, B, was calculated as Fitts throughput [2,28] for each successful trial whose starting position was not already inside the target on every 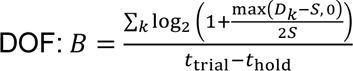. Here, D_k_ is the distance from the starting position to the target center on DOF k, S is the target radius, t_trial_ is the trial duration and t_hold_ is the hold time. Bend-sensor and Freebie runs were compared for hand control, and bend-KF and Freebie-KF runs for closed loop. One-and two-DOF runs were pooled for subsequent analyses.

### 2.10 Kalman filter decoding

#### Online decoders

Each closed-loop KF was trained on one hand-control run from the same day, using whichever signal (Freebie or bend sensor) had served as the label source during that run, and tested in a later run. Features were the mean absolute value (MAV) of the eight EMG channels in 32 ms bins. The kinematic labels for each bin were defined as the mean finger: position and a look ahead velocity (Δ mean finger position). The data were split chronologically 99%/1%, and the KF was fit as described below. Online, a BRAND node computed MAV over the most recent 32 ms of EMG every 32 ms and passed it to the KF. For Monkey N, closed-loop KFs were trained and run on the xPC system for four bend-sensor KF runs and four Freebie KF runs (three with index-angle labels and one with PIP y labels).

#### Offline decoders

Offline decoders were fit and validated within each hand-control run so that the two label sources could be compared on identical EMG data. For Monkey R, offline KFs were fit for every hand-control run (n = 24). For each run, two KFs were fit to identical EMG data, bins and time windows, one to the bend sensor and one to the Freebie signal, so that the label source was the only difference. EMG was first averaged as |x| in 4 ms bins, which yields the same 32 ms MAV as the raw samples, and analysis was limited to in-mode trials within the task window, skipping each run’s first trial.

For Monkey N (Fig. S4), KFs were trained offline on each of the 29 bend-sensor sessions that met the criteria in Section 2.5, with the finger-position labels set in turn to (i) the bend sensor and (ii) each Freebie index signal (index angle or PIP y), paired with MRS x in two-DOF sessions. All KFs of a session were trained and validated on the same frames.

#### Features

For experiments with Monkey R, EMG channels were binned into non-overlapping 32 ms bins, and the MAV of each channel was computed per bin. The EMG and Freebie kinematic streams were each was binned using their respective acquisition timestamps, without resampling the EMG. Bins spanned the union of EMG and kinematic time ranges, and a bin was dropped when either stream had no sample in it, which limited training to bins containing at least one Freebie frame. The kinematic state within each bin was defined as the mean finger position and the velocity (the next bins mean position minus the current bins mean position). For experiments with Monkey N, the real time xPC controller synchronously binned EMG and the most recently received Freebie kinematics on a shared experiment time.

#### Model

The KF state was [position, velocity, 1] per DOF. The observation matrix C was fitted by least squares, C = XᵀY(YᵀY)⁻¹, and the observation noise Q was the covariance of its residuals. The state-transition matrix A had identity on position, a unit position-velocity coupling term, a least-squares velocity-to-velocity block and a constant bias. The process noise W was the residual covariance of A, with its position and bias rows and columns set to zero. Offline prediction started from a zero state with error covariance W.

#### Evaluation

The binned data were split chronologically into the first 80% for training and the last 20% for validation, without shuffling. Validation performance was reported for the position outputs as the Pearson correlation and the normalized mean-squared error (MSE divided by the variance of the validation target). Monkey R performed only a one-DOF task, so each KF decoded index finger alone. Monkey N performed both one-DOF and two-DOF sessions: in one-DOF sessions the KF decoded index position only, and in two-DOF sessions it decoded index and MRS positions together. Index finger results combine KFs from both session types. MRS results come only from the two-DOF sessions. While in the Freebie condition, they are taken from the KF whose labels were the index angle and MRS x coordinate.

### 2.11 Statistical analysis

All tests were two-sided with α = 0.05 and used SciPy 1.17.1. Independent groups were compared with the Mann-Whitney U test on run means, one value per run, bend sensor versus Freebie hand-control runs, and bend-KF versus Freebie-KF closed-loop runs. Paired comparisons used the Wilcoxon signed-rank test, matched by run: offline-KF metrics between label sources and, for Monkey N, |r| between Freebie features. Multiple comparisons: within each figure panel all pairwise comparisons were tested, and p-values were corrected with the Holm-Bonferroni method across the comparisons in that panel.

### 2.12 Software and data availability

Analyses used Python with NumPy 2.4.4, pandas 2.3.3, SciPy 1.17.1, PyNWB 3.1.3 and Matplotlib 3.10.9, and our lab’s *neuraldecoding* package for NWB loading, online metrics, feature extraction and the Kalman filter. Pose estimation used DLC 3.0.0rc10 with PyTorch 2.8.0. Real-time experiments with Monkey R ran with BRAND [25], and human 3D tracking used MediaPipe Hand Landmarker [13] and aniposelib [17]. The code to run Freebie during live experiments is available at github.com/draeloslab/improv/tree/Freebie/experiments/Freebie. Freebie paper code, including figure generation code, is available at github.com/draeloslab/joseph_2026. The data supporting the conclusions of this article will be made available by the authors upon reasonable request.

## 3. Results

Freebie is a markerless, camera-based system that measures hand and finger positions in real time. We evaluated whether it could replace an instrumented manipulandum both as a direct control signal and as a source of labels for a real-time, closed-loop EMG decoder. Monkey R performed a one-DOF index finger target acquisition task while bend sensor signals and implanted forearm EMG were recorded simultaneously. One industrial camera viewed the index finger, a DLC network estimated the PIP joint location, and the Kalman-filtered keypoint was streamed over UDP to a BRAND real-time computer, which mapped it onto finger position (Fig. 1). We also present data from Monkey N, recorded with an earlier two-camera version of the system that communicated with an xPC computer over a serial link.

### 3.1 Freebie closely tracked static finger joints

In a motor BMI, every subsequent processing step depends on the accuracy of the step before it, starting with the pose estimation network. We first assessed agreement between network predictions and manual annotations. Predictions were compared with manual labels on held-out frames sampled from experimental session videos (Fig. 2A). Only predictions with likelihoods ≥ 0.99 were included, and errors were referenced to 20 px, the image diameter of the smallest labeled finger width. For the top camera, median Euclidean errors were 2.35 px for PIP, 3.79 px for DIP, 1.68 px for MCP and 6.54 px for the wrist (n = 80, 68, 59 and 15 frames, respectively). The median error for the MRS keypoint in the side camera view was 5.57 px (n = 23 frames). The maximum error for any keypoint was 14.3 px, so every error was under the 20 px benchmark (Fig. 2B). Thus, static tracking accuracy was unlikely to limit downstream system performance.

### 3.2 Freebie operated within the real-time frame budget

A tracking system can support closed-loop control only if it produces an estimate for the current frame before the next frame arrives. We therefore measured the interval from the camera callback to UDP transmission of pose estimates for every frame across 32 runs (690,485 frames) and quantified the contribution of each processing stage. Median end-to-end latency was 15.8 ms in the 14 runs recorded at 30 FPS (83,166 frames) and 16.2 ms in the 18 runs recorded at 60 FPS (607,319 frames) (Fig. 3A). Inference accounted for most of the processing time (14.6 ms, mean of run means), followed by frame acquisition (1.3 ms) and the sender (0.12 ms) (Fig. 3B). Latency was stable across runs, with run-mean end-to-end latencies between 15.7 and 16.5 ms.

**Figure 3.**
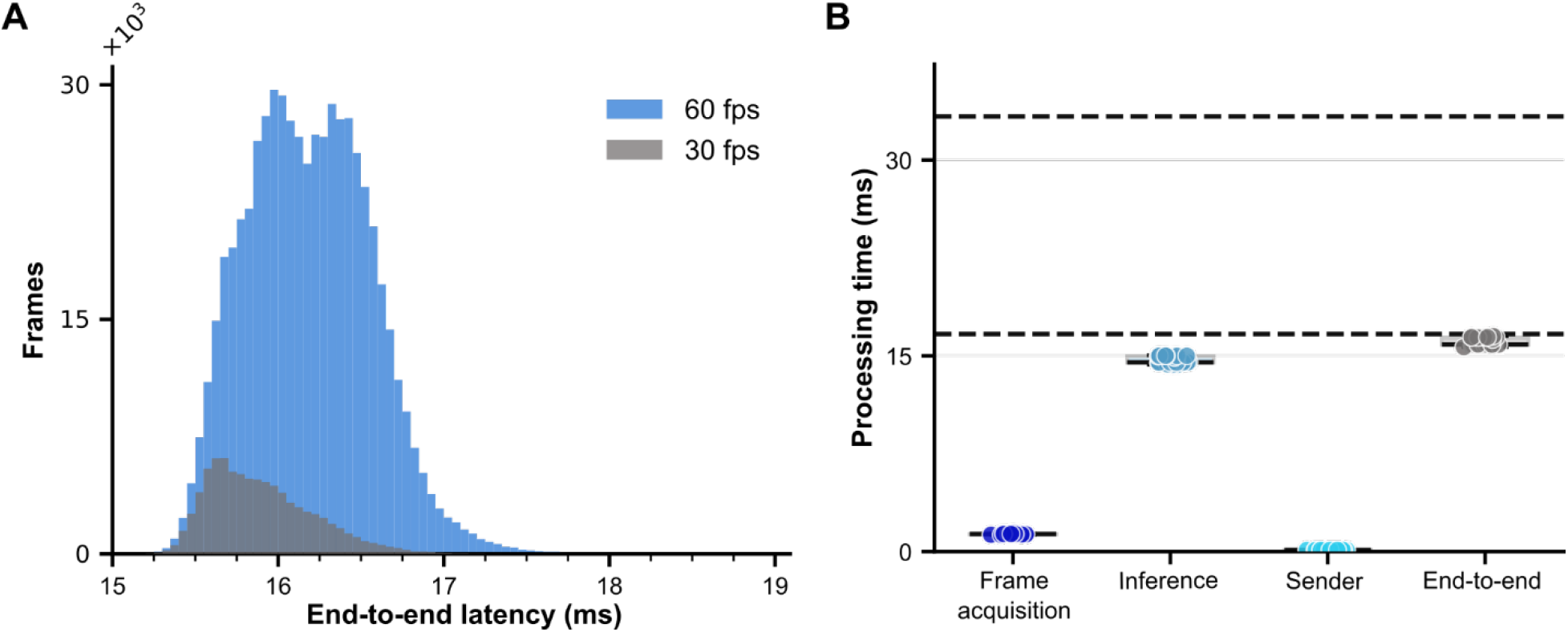
Freebie operated within the real-time frame budget. (**A**) Distribution of end-to-end latencies, from the camera callback to UDP transmission, for all frames from runs recorded at 30 FPS (grey; n = 14 runs, 83,166 frames) and 60 FPS (blue; n = 18 runs, 607,319 frames). (**B**) Processing time of each pipeline stage (frame acquisition, inference, sender) and end-to-end, with one point per run mean (n = 32 runs). Dashed lines mark the 30 and 60 FPS frame budgets (33.3 and 16.7 ms).

Because closed-loop control depends on the upper tail of the latency distribution rather than its median alone, we also examined frames that exceeded the frame budget. At 30 FPS, only one of 83,166 frames (0.001%) exceeded the 33.3 ms budget (maximum 43.0 ms). At 60 FPS, the median latency of 16.2 ms was within the 16.7 ms budget on average, though 9.9% of frames exceeded it. Late frames were still delivered, and frames were rarely lost: across all runs, the processor skipped 0.012% of frames (5 of 83,166 at 30 FPS; 75 of 607,319 at 60 FPS) and the camera failed to deliver 0.013% of frames (7 and 81 frames, respectively). These losses occurred in short bursts (longest: 6 skipped and 12 missing frames, ≤200 ms at 60 FPS). With the earlier version of Freebie, median end-to-end latency was 28.3 ms, and 12.8% of its frames exceeded the 33.3 ms budget (Fig. S2). Together, these benchmarks demonstrate that Freebie reliably delivered kinematic estimates well within the tens of milliseconds latency budget and small frame loss tolerances required for responsive, closed loop neuroprosthetic control.

### 3.3 Freebie signals closely tracked bend sensor measurements

Before evaluating whether Freebie could replace the bend sensors, we assessed whether the two systems measured comparable finger movements. In Monkey R’s hand-control runs, Freebie’s PIP y signal closely tracked the simultaneously acquired bend sensor signal measuring index flexion (median |r| = 0.96, interquartile range 0.95–0.97, n = 24 runs; Fig. 4A, B). In Monkey N, we evaluated three finger movements metrics and found that their agreement with bend sensors depended on the Freebie feature: PIP y and MRS x correlated well with their corresponding bend sensor signals (median |r| = 0.86 and 0.84, n = 30 and 16 sessions), whereas the derived three-point index joint angle did not (median |r| = 0.41, n = 30; Wilcoxon signed-rank, Holm-corrected p < 0.001 versus PIP y; Supplementary Fig. S3A, B). Raw keypoint coordinates therefore provided more reliable kinematic signals than the derived joint angle, and our final system used the PIP y coordinate throughout all runs.

**Figure 4.**
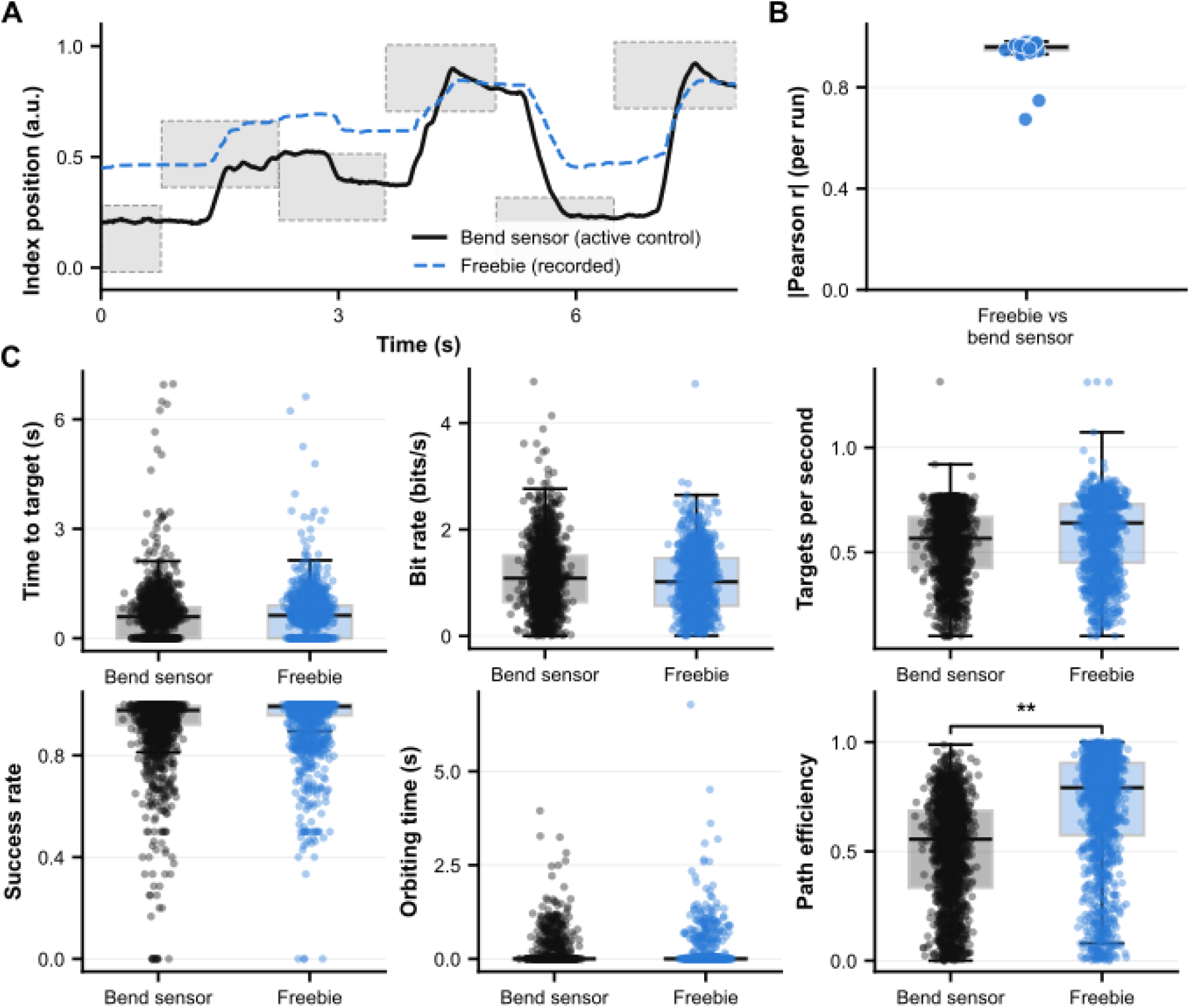
Direct Freebie control matched bend sensor control. (**A**) Example trials in which the bend sensor (black) drove the virtual hand while Freebie (blue, dashed) was recorded concurrently. Grey boxes are target zones. (**B**) Absolute Pearson correlation between Freebie and the bend sensor, one point per hand-control run (n = 24). (**C**) Online performance under bend sensor (15 runs) and Freebie (9 runs) control: success rate, bit rate, targets per second, time to target, orbiting time and path efficiency. Dots are trials; success rate and targets per second are running values within each run. Time to target, orbiting time and path efficiency are computed on successful trials. Statistics were computed on run means (Mann-Whitney U; **p < 0.01).

### 3.4 Direct Freebie control matched bend sensor control

To assess whether Freebie could directly replace the bend sensor during online hand control, the virtual hand source was switched from the bend sensor to Freebie, with all other task parameters unchanged. Freebie control performed as well as bend sensor control (n = 9 Freebie and 15 bend sensor runs; 1,600 and 1,796 trials; Fig. 4C). Freebie and bend sensor control did not significantly differ in success rate (median of run means, 0.97 vs 0.99; Mann-Whitney U, p = 0.29), bit rate (1.02 vs 1.06 bits s⁻¹, p = 0.21), targets per second (0.50 vs 0.57 targets s⁻¹, p = 0.63), time to target (0.66 vs 0.72 s, p = 0.37) or orbiting time (0.11 vs 0.13 s, p = 0.37). Path efficiency was the only metric that differed significantly between control modalities and was higher under Freebie control (0.67 vs 0.52, p = 0.005; Fig. 4C).

With an earlier pipeline, Freebie control driven by the index joint angle underperformed bend sensor control on most metrics (median success rate 0.29 vs 0.94; Holm-corrected p < 0.001; Fig. S3C). The two runs driven by PIP y, however, did not differ significantly from bend sensor control except in time to target (1.00 vs 0.50 s, p = 0.003). The comparable performance between Freebie and bend sensors in trials with Monkey R is consistent with Freebie’s low latency, lighter smoothing than the 10-frame moving average used earlier for Monkey N, and the use of the PIP y coordinate throughout.

### 3.5 Freebie-and bend-sensor-trained decoders achieved comparable closed-loop control

Kinematic labels are used to train BMI decoders, so we evaluated whether Freebie could replace the bend sensor as the source of training labels. Monkey R controlled the task using a KF trained on Freebie or bend sensor signals (n = 4 runs per group; Fig. 5A). Performance was not significantly different between decoder conditions. For bend sensor and Freebie-trained decoders respectively, success rates were 0.77 and 0.98, bit rates were 1.77 and 1.56 bits s⁻¹, targets per second was 0.49 vs 0.45 targets s⁻¹, time to target was 0.65 and 0.87 s, orbiting time was 0.21 and 0.23 s, and path efficiency was 0.72 and 0.68 (all Mann-Whitney U; p > 0.5; Fig. 5B). Monkey N also demonstrated closed loop control with decoders trained on both conditions for the 2DOF task (Fig. S4A, B), and performance likewise did not differ between bend-sensor-trained (4 sessions) and Freebie-trained (3 sessions) decoders (Fig. S4C). Thus, decoders trained on Freebie-derived labels supported closed-loop control and were comparable to decoders trained on bend sensor labels.

**Figure 5.**
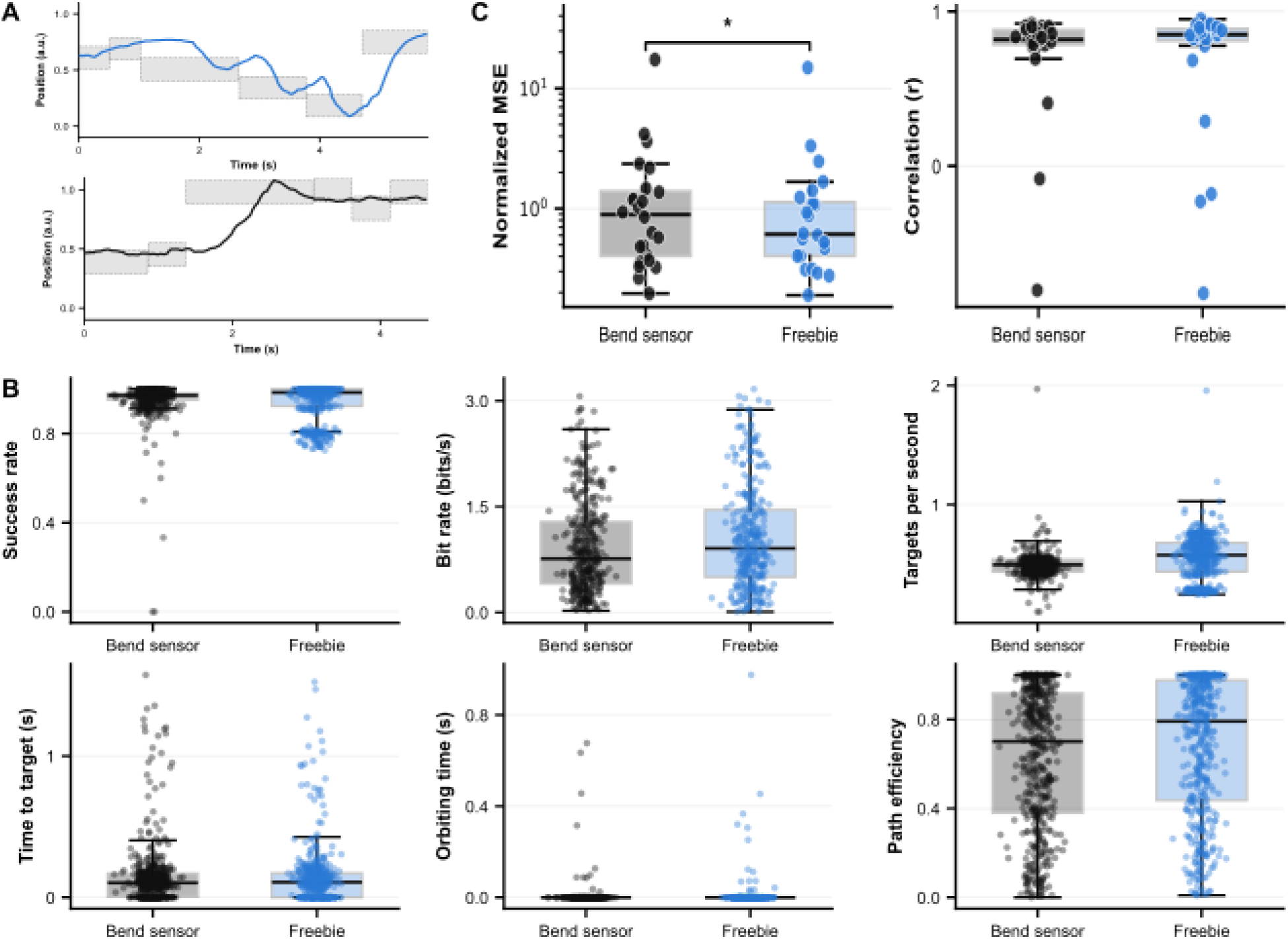
Kalman filters trained on Freebie-derived labels support closed-loop control. (**A**) (Top) Example closed-loop decoded index position from a KF trained on Freebie labels and (Bottom Grey boxes are target zones.) for a KF trained on bend sensor labels.(**B**) Online closed-loop performance of KFs trained on bend sensor and Freebie labels (4 runs each): success rate, bit rate, targets per second, time to target, orbiting time and path efficiency. Dots are trials; statistics on run means (Mann-Whitney U). (**C**) Offline KF performance for the index DOF, one point per hand-control run (n = 24), with the two label sources fit to the same EMG data. Left: normalized MSE (MSE divided by the variance of the validation target; log scale). Right: correlation between decoded and target positions (Wilcoxon signed-rank tests. *p < 0.05).

To determine whether label source affected how well the decoder learned the mapping from EMG activity to finger position, we also compared the two label sources offline (Fig. 5C). For each of Monkey R’s 24 hand-control runs, two KFs were fit to identical EMG data, one with bend sensor labels and one with Freebie labels. Freebie-trained KFs had lower normalized MSE on held-out data (median 0.61 vs 0.89; Wilcoxon signed-rank, p = 0.016) and similar correlation with the target signal (median r = 0.85 vs 0.82, p = 0.39). Some runs decoded poorly with either label source: normalized MSE exceeded 1 in at least a quarter of runs, and two runs per label source had negative correlations (Fig. 5C). This suggests that run-specific factors, rather than label source, limited decoding in those runs. In Monkey N, KFs trained on labels from the earlier pipeline were less accurate offline than bend-sensor-trained KFs (median index correlation 0.59 for the index angle and 0.62 for PIP y vs 0.88; n = 29 sessions; Holm-corrected p < 0.001; Fig. S4D), yet this offline deficit did not produce detectably worse closed-loop performance in the small number of runs tested (Fig. S4C).

### 3.6 Real-time tracking of free behavior and stimulated movements

We have also extended Freebie beyond manipulandum-based tracking. Monkey R performed the same one DOF task as above but with the manipulandum removed and the virtual hand driven by Freebie alone (Supplementary Fig. S5A, Video 1). Freebie control without the manipulandum performed similarly to bend sensor control with it: 98.9% and 100% of trials in the two runs, with a median bit rate of 1.05 bits/s, against 100% and 1.00 bits/s with the bend sensor (Fig. S5B).

Next, we tracked 21 keypoints per hand with MediaPipe [13] during a free behavior task (Fig. 6A), in which the monkey reached for and interacted with a toy by pinching and spinning several objects. Five cameras with overlapping fields of view recorded the hand concurrently, and Freebie tracked the hand in 2D in each view in real time. A manipulandum cannot measure such unconstrained, higher dimensional hand which motivates replacing it. Manipulandum experiments showed that a single view reliably captures only one DOF. For example, the top view camera could not resolve MRS movement. The side camera could see MRS but not the wrist, so an index MCP angle could not be computed from it. Two DOF control therefore required a second, independently running side camera. Our free behavior task also presented this same challenge. For example, treating each camera independently, in one session, the index fingertip was detected in 6.7% to 73.1% of frames depending on the view. Most missed detections occurred when the monkey’s fingers were curled into the toy, hiding the fingertips behind the toy and the back of the hand (gaps in Fig. 6A, bottom). By combining the views at each step, at least one camera detected the fingertip in 84.7% of images. Multi-camera 3D tracking is designed to resolve these occlusions, because a keypoint hidden in one view is often visible in another.

**Figure 6.**
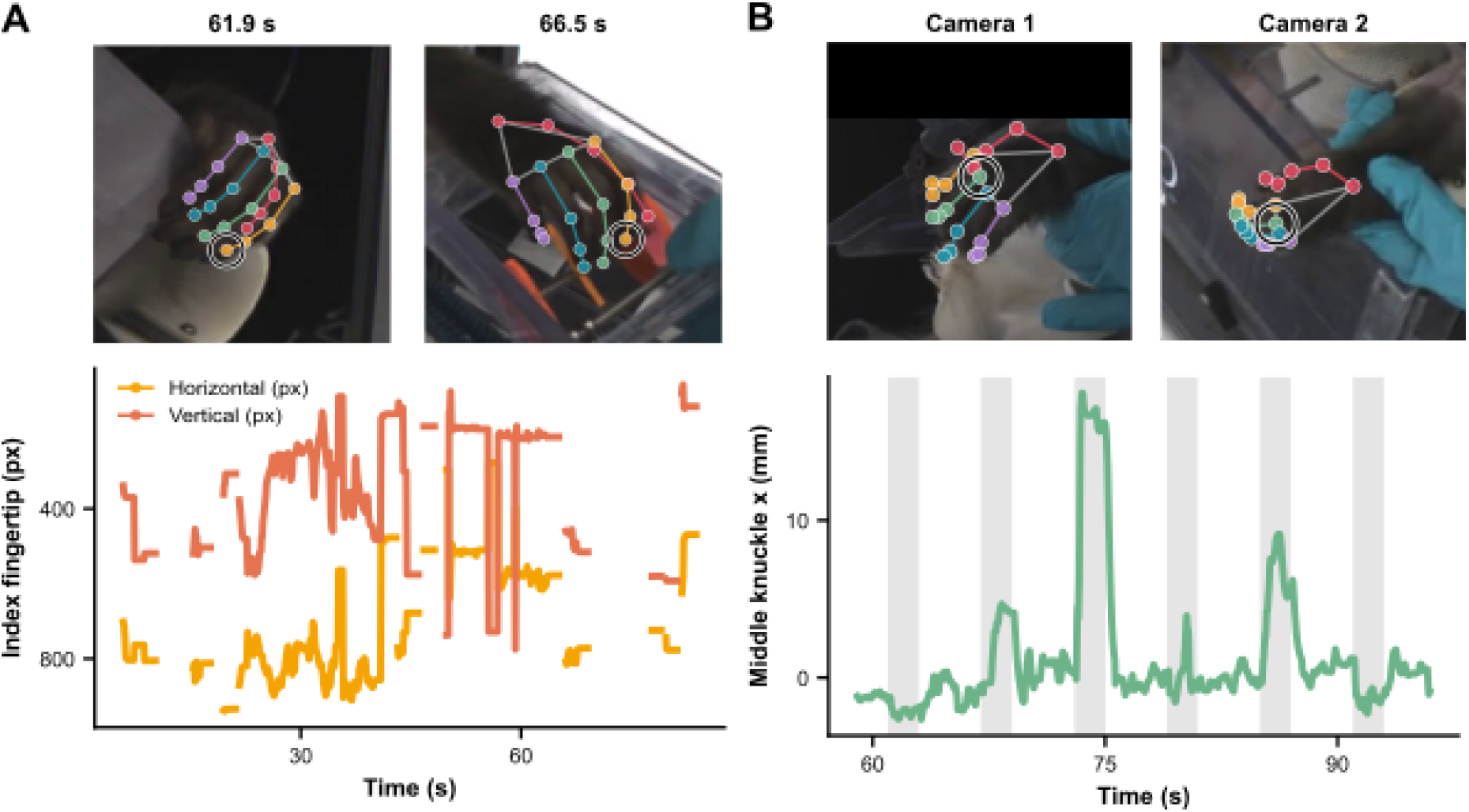
Extension of Freebie beyond manipulandum tasks. (**A**) Real-time tracking of 21 hand keypoints during a free behavior session. Top: example frames (acquired at 61.9 and 66.5 s) with predicted keypoints and skeleton overlaid, one color per digit. Bottom: index fingertip position in one camera view (x, dark; y, orange; pixels). Gaps are periods in which the fingertip was not detected. (**B**) Pseudo-live tracking during an automated FES characterization experiment. Top: simultaneous views from two cameras with predicted keypoints. Bottom: x position (mm) of the middle-finger MCP joint across different stimulations reconstructed in 3D from 2 cameras. Shaded bars mark stimulation periods.

Lastly, we used Freebie to measure joint positions during automated characterization of functional electrical stimulation (Fig. 6B). Stimulation trains with randomly selected 2-3 multi electrode combinations at a constant current and pulse width with a stim and rest period of 2 and 4 seconds, were delivered automatically while two cameras recorded the hand, and keypoints were triangulated to 3D; tracking was run in pseudo-live mode (recorded video passed through the real-time pipeline as if live). Different stimulation trains evoked clearly different movements of the middle-finger MCP joint, ranging from almost no displacement to extensions of roughly 18 mm (Fig. 6B, bottom), showing that Freebie can resolve stimulation-specific joint movements in physical units. Freebie is particularly useful in this setting because the hand must remain free to move in response to stimulation. Real-time kinematic feedback supplied by Freebie could also support adaptive optimization of stimulation parameters for improving FES control.

## 4. Discussion

Advancing intracortical neuroprosthetics towards naturalistic, multi-finger dexterity requires moving beyond the physical constraints of instrumented manipulanda. Here, we developed Freebie, the first real time, multi camera markerless hand tracking platform purpose built for closed loop primate neuroprosthetics and validated it across two monkeys both as a direct real-time control signal and as a source of training labels for neural decoding. Four principal findings demonstrate the system’s current capabilities. First, Freebie delivered keypoint estimates with a median end-to-end latency of ∼16 ms at both 30 and 60 FPS. Only one frame exceeded the frame budget at 30 FPS, and 9.9% of frames when acquiring images at 60 FPS. Additionally, about 0.01% of frames were skipped by the processor or never delivered by the camera. Second, Freebie-derived signals closely agreed with bend sensor measurements. Third, direct Freebie control performed as well as bend sensor control. Fourth, decoders trained on Freebie-derived labels performed similarly to decoders trained on bend sensor labels, both online and offline. Freebie also supported direct hand control with the manipulandum removed. Because neither direct control nor decoder training requires an instrumented manipulandum, Freebie can enable kinematic labeling of movements and postures that a manipulandum cannot readily measure, including unconstrained and higher-dimensional hand movements.

The comparison between the two monkeys indicates several design choices that may influence direct control performance. With Monkey N’s earlier pipeline, Freebie control underperformed bend sensor control, primarily because of longer times to target and orbiting times, though path efficiency was unaffected (Supplementary Fig. S3C). This pattern is consistent with a delayed rather than spatially noisy control signal. In that pipeline, a 10-frame moving average at 30 FPS added a group delay of (k − 1)/2 = 4.5 frames (∼150 ms) to the 28.3 ms processing latency, so the virtual hand lagged by roughly 180 ms. Control-loop delays of 100–300 ms measurably degrade BMI performance, producing slower, less direct and less smooth movements [29]. In the current system, processing latency was 16 ms and the keypoint was smoothed with a light exponential moving average. This added a group delay of about 7 ms at 60 FPS (14 ms at 30 FPS), for a total lag of only 23 ms at 60 FPS (30 ms at 30 FPS), compared with ∼180 ms of lag for the original system. Moreover, the control signal was usually the index joint angle, which agreed poorly with the bend sensor value (median |r| = 0.41 vs 0.86 for PIP y). Using the PIP y coordinate throughout removed the performance deficit (Fig. S3B). Because these two changes were made together, and the two monkeys also differed in task and rig, we cannot attribute the final improvement to a single factor. However, the two Monkey N runs driven by PIP y still had longer times to target than bend sensor control (Supplementary Fig. S3C), which suggests that delay contributed beyond the choice of feature.

Path efficiency was higher under Freebie control than under bend sensor control in Monkey R. However, path efficiency is calculated from the summed sample-to-sample path length and is therefore sensitive to noise and smoothing. The two control signals were conditioned differently: the bend sensor was averaged over 100 ms (20 samples at 200 Hz), adding a group delay of about 48 ms, whereas Freebie had a group delay of only 7 ms at 60 FPS (14 ms at 30 FPS). The observed difference in path efficiency may therefore reflect signal conditioning rather than better control with Freebie. Task settings, including target radius, hold time, and smoothing, also varied between runs and were pooled.

The difference between offline and online performance further demonstrates the practical utility of Freebie. Decoders trained with Freebie labels performed comparably online to bend sensor-trained decoders in both monkeys, though their offline results differed. In Monkey R, Freebie labels produced lower normalized error and similar correlation. In Monkey N, decoders with an earlier Freebie model showed lower offline accuracy (index correlation 0.62 with PIP y and 0.59 with the index angle, versus 0.88 with the bend sensor) but did not ultimately impair closed-loop control (Supplementary Fig. S4C). Monkey N’s Freebie labels were likely noisier: the index angle tracked the bend sensor poorly, and signals were rescaled using post-hoc percentiles rather than calibrated ranges. Monkey R’s PIP y signal tracked the bend sensor closely and was calibrated to a fixed pixel range in each run. Overall, our results nevertheless show that an offline deficit need not translate into impaired closed-loop control. Within the limits of this study, bend sensors therefore appear replaceable where it matters most: markerless keypoints alone provided the labels needed to train a usable EMG decoder without instrumenting or constraining the hand.

Existing real-time tools, including DLC-Live and SLEAP, report single-frame inference latencies comparable to or lower than Freebie’s [15,23]. The main difference, however, lies in how the resulting features are integrated into the experimental system. Previous methods have primarily used a single camera to trigger discrete contingent events, such as the illumination of an LED or delivery of optogenetic stimulation, for which only the timing of detection matters. In contrast, Freebie continually streams position estimates to a real-time experimental controller as a replacement for an instrumented kinematic sensor and has been integrated with two different real-time systems (xPC Target and BRAND). Freebie also supports multiple camera views, demonstrated here with real-time one and two camera tracking in monkey experiments as well as four camera tracking for 2-4 DOF closed loop control by a human (Fig. S1B). Adding more DOFs to the task did not meaningfully increase system latency (median 12.2–12.8 ms for two to four DOFs), because every joint angle is computed from the same 21 tracked keypoints (Fig. S1C).

This study had several limitations. First, the closed-loop decoder comparison included only four runs per condition. Monkey N provides additional evidence, but it was recorded with a different version of the system, task, and rig. Second, the study evaluated a limited set of somewhat constrained movements, which restricts the generalizability of our results to higher-dimensional or more naturalistic behaviors. Third, all closed-loop decoders mapped forearm EMG activity, rather than intracortical activity, to finger position. EMG served here as an accessible proxy for a neural control signal. Because a decoder learns a mapping from its input signal to whatever labels it is given, we expect the label-quality result to extend to intracortical decoders. However, EMG is more directly coupled to finger movement than cortical activity is, and the sensitivity of intracortical decoders to label noise remains to be tested.

Several measurement limitations also apply. Although the cameras were rigidly mounted and calibrated once, we did not quantify calibration drift across sessions. End-to-end latency was measured from the camera callback to packet transmission, an interval that excludes sensor exposure, readout and transfer before the callback, and the interval between the experimental computer and the rendered display. The total delay experienced by the monkey thus likely exceeded the reported median. In Monkey N, the DIP and PIP joints were occluded by the manipulandum, and their annotations therefore represented inferred rather than directly observed joint locations. Ink marks were applied to selected joints and we did not test whether model performance depended on these marks. Finally, bend sensor and Freebie signals were aligned using xPC timestamps under the assumption of constant transport delay, which may have resulted in conservative estimates of agreement.

## 5. Conclusion

To our knowledge, Freebie is the first markerless pose estimation platform to integrate synchronized multi camera acquisition, real time 3D kinematic reconstruction, and bidirectional hardware communication within the strict ∼16 ms latency budget required for closed loop primate neuroprosthetics. With our current pipeline, Freebie served as an effective direct control signal for hand and finger tasks, matching the performance of tasks using bend sensor control. Its keypoints successfully served as training labels for EMG decoders that performed comparably to decoders trained on bend sensor labels, both offline and in a small number of closed-loop runs. An earlier version of the pipeline with slightly higher latency and a less reliable joint-angle feature still produced usable decoder training labels despite lower offline accuracy. These findings indicate that markerless keypoints can replace manipulandum measurements both for direct hand control and as decoder training labels, and that offline decoding metrics may underestimate the online utility of markerless labels. We also demonstrated Freebie in experiments beyond the manipulandum: it supported direct hand control after the manipulandum was removed, tracked 21 hand keypoints during unconstrained behavior and measured stimulation-evoked joint movements in 3D during automated FES characterization. By replacing restrictive mechanical sensors with real-time multi-view image analysis, Freebie provides the foundational infrastructure needed to train intracortical decoders and calibrate multi-muscle FES systems during natural, unconstrained hand use.

## Acknowledgments

Thank you to Eric Kennedy for his animal and experimental support, the University of Michigan Unit for Laboratory Animal Medicine for their expert veterinary and surgical support, and the University of Michigan Biointerfaces Institute for their support.

Primary funding to support this project was provided by the National Institutes of Health under award numbers R01NS134834 for J.D., M.M., A.W., and T.K.; and R01DA063389 for A.D. J.J. had support from the National Institutes of General Medical Sciences training grant T32GM141746 and the University of Micham Rackham Merit Fellowship. M.C. had support from the European Union’s NextGenerationProgramEU award MNESYS PE0000006. L.L. acknowledges support from a Magnificent Michigan Summer Undergraduate Research Fellowship. M.K., H.T., and M.G.C. had funding from the National Science Foundation’s EFRI BRAID award 2223822. M.M. had support from the National Science Foundation award 1926576. O.L. was supported by the National Science Foundation Graduate Research Fellowship program award number DGE-2241144. M.K. and C.C. are supported by the Michigan Biosciences Faculty Award Recognition. A.D. is supported as an Alfred P. Sloan Research Fellow in Neuroscience (FG-2024-21442), and by a Career Award at the Scientific Interface from the Burroughs Welcome Fund (1021865.01).

## Supplementary Material

**Fig. S1:**
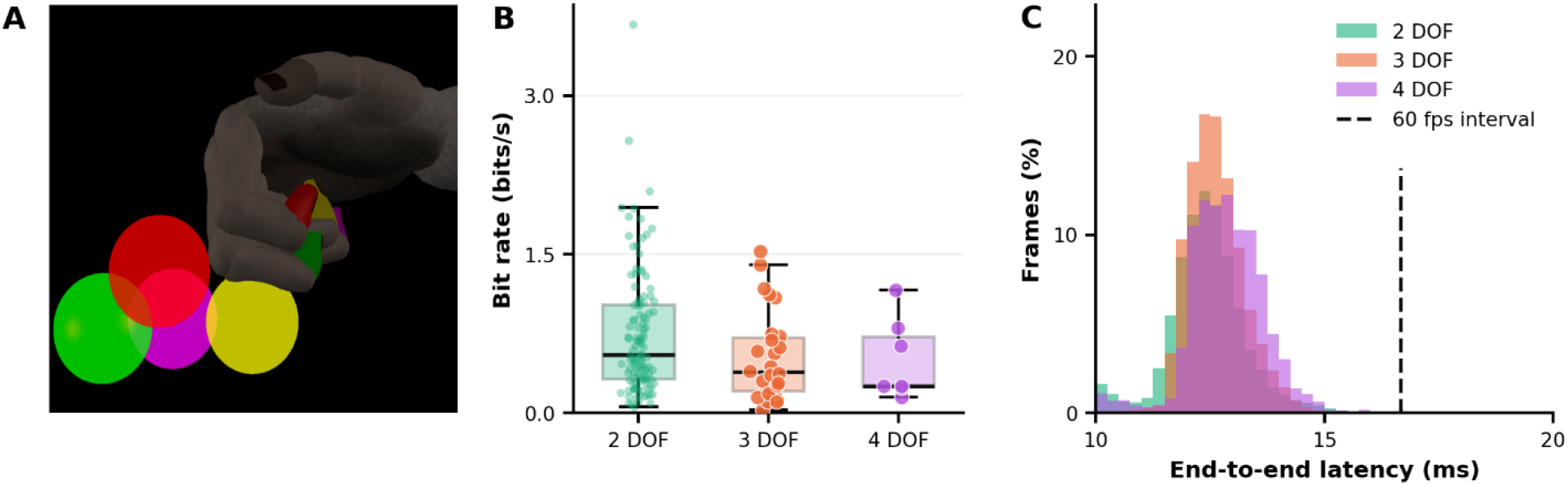
Real-time 3D hand tracking enabled 2–4 degree-of-freedom control for human hands. **(A)** The task screen. A virtual hand, driven by tracked human finger angles, moves toward spherical targets. This example is the four-DOF configuration. **(B)** Bit rate of individual successful trials for two (n = 115 trials), three (n = 26) and four (n = 7) DOF, one run per DOF count (success rates 88%, 69% and 50% of 154, 45 and 14 trials). Medians are 0.55, 0.39 and 0.26 bits per second. **(C)** End-to-end latency, as a percentage of frames, for two (n = 24,018 frames), three (n = 12,515) and four (n = 2,234) DOF. Each pose was triangulated from four cameras running at 60, and latency is measured from the newest of the four frames contributing to that pose (median 12.2, 12.6 and 12.8 ms for two, three and four DOF). Dashed line marks 60 FPS budget (16.7 ms).

**Figure S2.**
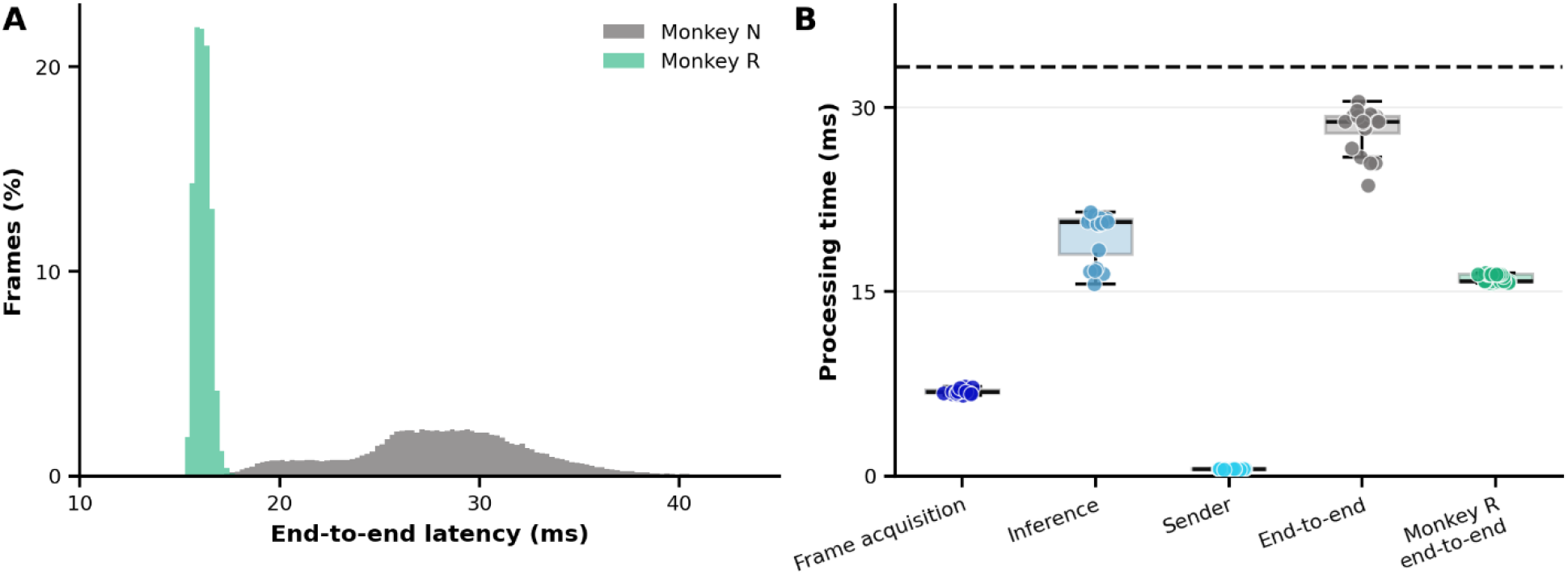
Latency of the earlier pipeline (Monkey N) compared with the current pipeline (Monkey R). (**A**) Distribution of end-to-end latency for frames pooled across Monkey N’s sessions (grey; n = 20 sessions, 144,888 frames; median 28.3 ms) and Monkey R’s runs (green; n = 32 runs, 406,873 frames; median 15.9 ms). (**B**) Processing time of each pipeline stage (frame acquisition, inference, sender) and end-to-end for Monkey N, one point per session mean (n = 20), and Monkey R’s end-to-end latency, one point per run mean (n = 24). The dashed line is the 30 FPS frame budget (33.3 ms).

**Figure S3:**
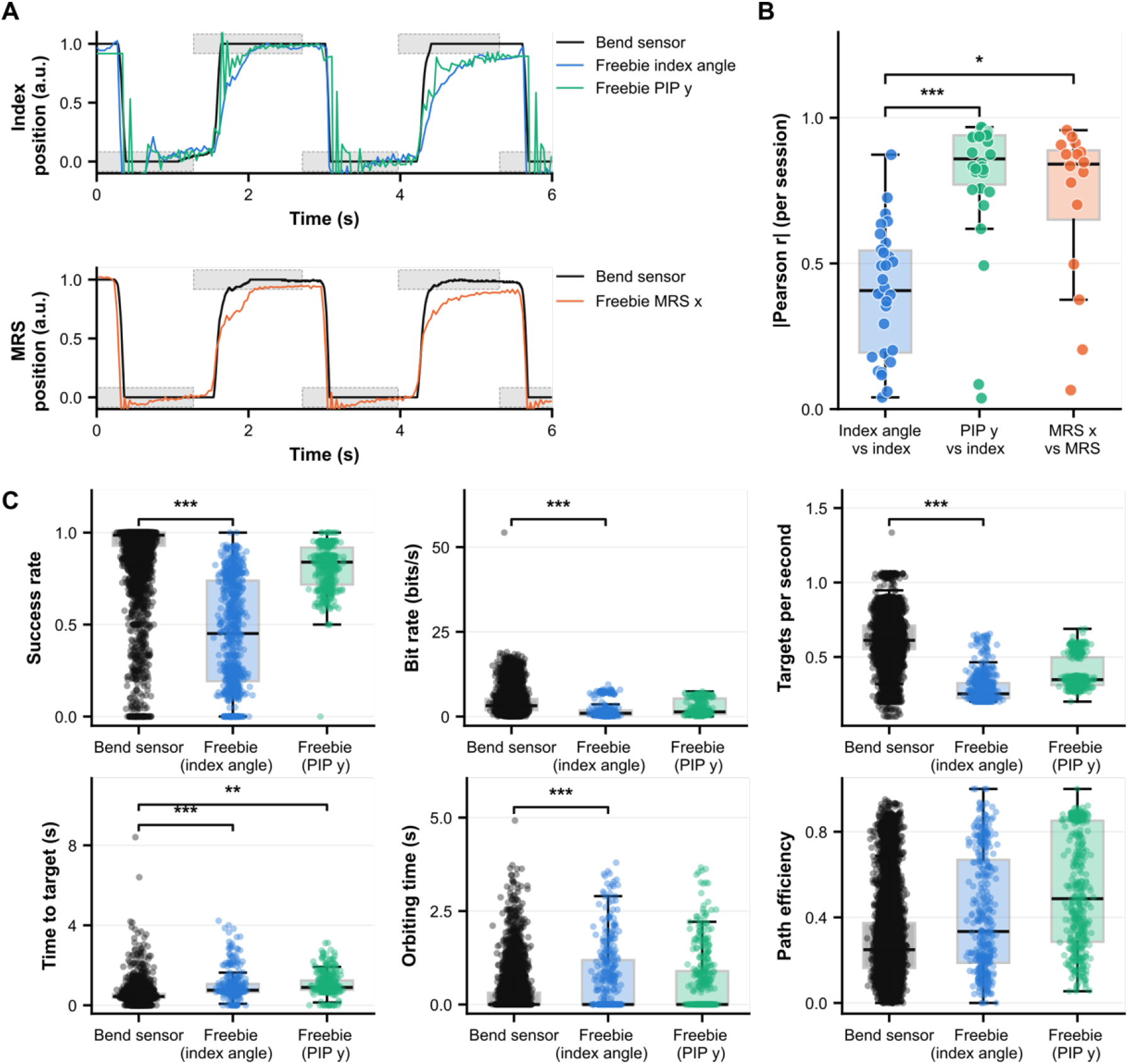
Real-time hand control with Freebie-derived features and bend sensor signals. (**A**) Example trials in which the bend sensors controlled the virtual hand while Freebie operated concurrently. Top: index finger position measured by the bend sensor (black), and Freebie-derived index angle (blue) and PIP y coordinate (green). Bottom: MRS position measured by the bend sensor (black) and the Freebie MRS x coordinate (orange). Signals were scaled to 0–1; shaded grey regions indicate target zones. (**B**) Absolute Pearson correlation between each Freebie-derived feature and its corresponding bend sensor signal, with one point per session (index angle and PIP y coordinate n = 30; MRS x coordinate n = 16). Paired Wilcoxon signed-rank tests, Holm-corrected. (**C**) Online performance under bend sensor control and Freebie control, with Freebie runs split by the driving signal (index angle or PIP y): success rate, bit rate, targets per second, time to target, orbiting time and path efficiency. One point per trial; success rate and targets per second are running cumulative values within each run. Time to target, orbiting time and path efficiency are computed on successful trials only. Mann-Whitney U tests, Holm-corrected on session means with n=47-50 bend sensor, 13-15 index angle, and 2 PIP y sessions. *p < 0.05, **p < 0.01, ***p < 0.001.

**Figure S4.**
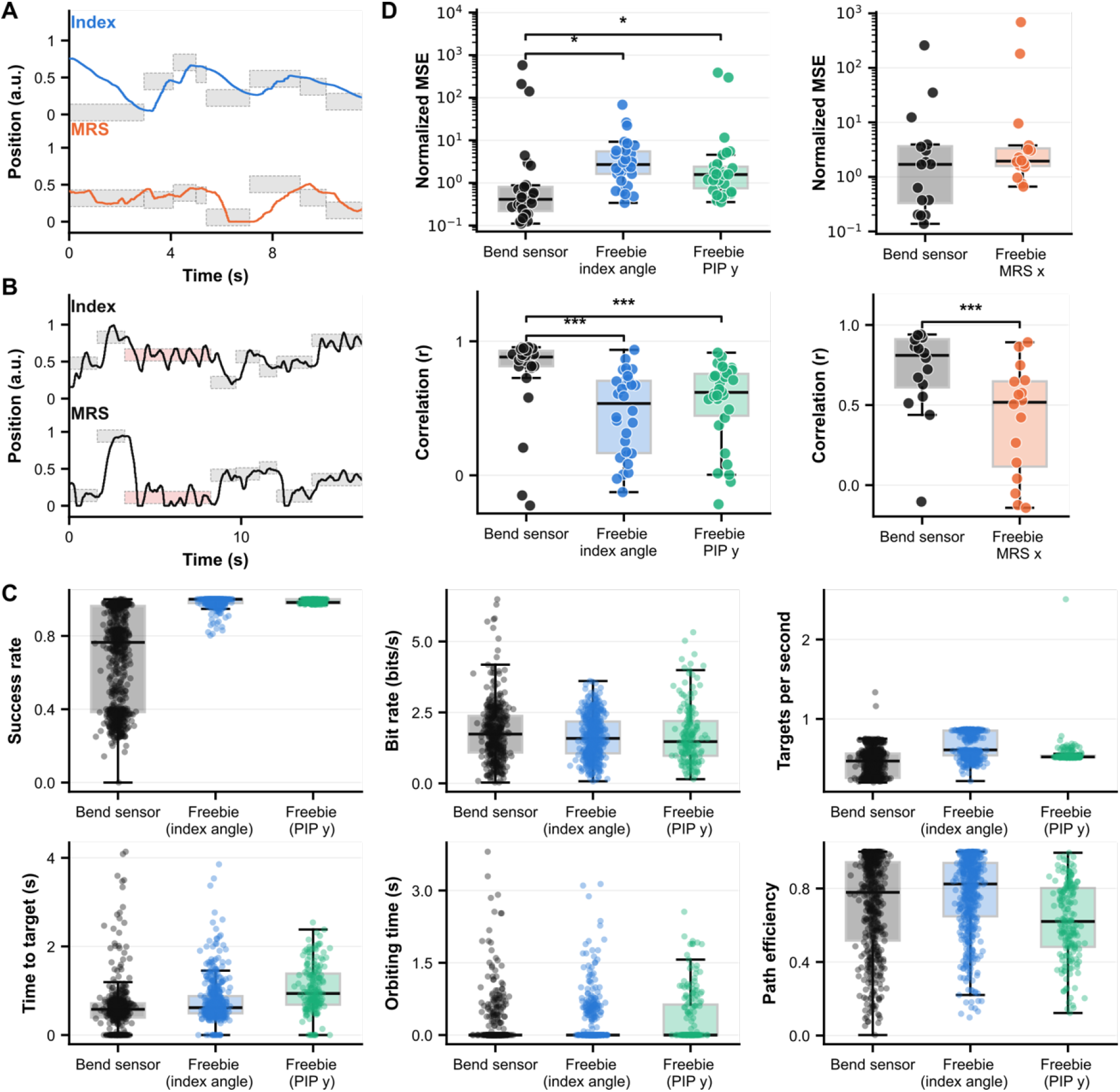
Kalman filters trained on Freebie-derived labels support closed-loop control. (**A**) Example closed-loop decoder output for the index (left) and MRS (right) DOFs from a KF trained on Freebie-derived labels. Grey boxes are target zones of successful trials. (**B**) Same as (A) for a KF trained on bend sensor labels. Red boxes are missed trials. (**C**) Online closed-loop performance of KFs trained on bend sensor labels and on Freebie-derived labels (index angle or PIP y): success rate, bit rate, targets per second, time to target, orbiting time and path efficiency. One point per trial; metrics are defined as in figure 4C. Mann-Whitney U tests on session means, bend sensor (4 sessions) versus index angle (3 sessions); the single PIP y run was not tested. (**D**) Offline KF performance for the index and MRS DOFs, one point per session. Decoders were trained on the first 80% of each session and evaluated on the final 20%. Left: normalized MSE (MSE divided by the variance of the validation target; log scale). Right: correlation between decoded and measured positions. Paired Wilcoxon signed-rank tests, Holm-corrected.

**Figure S5.**
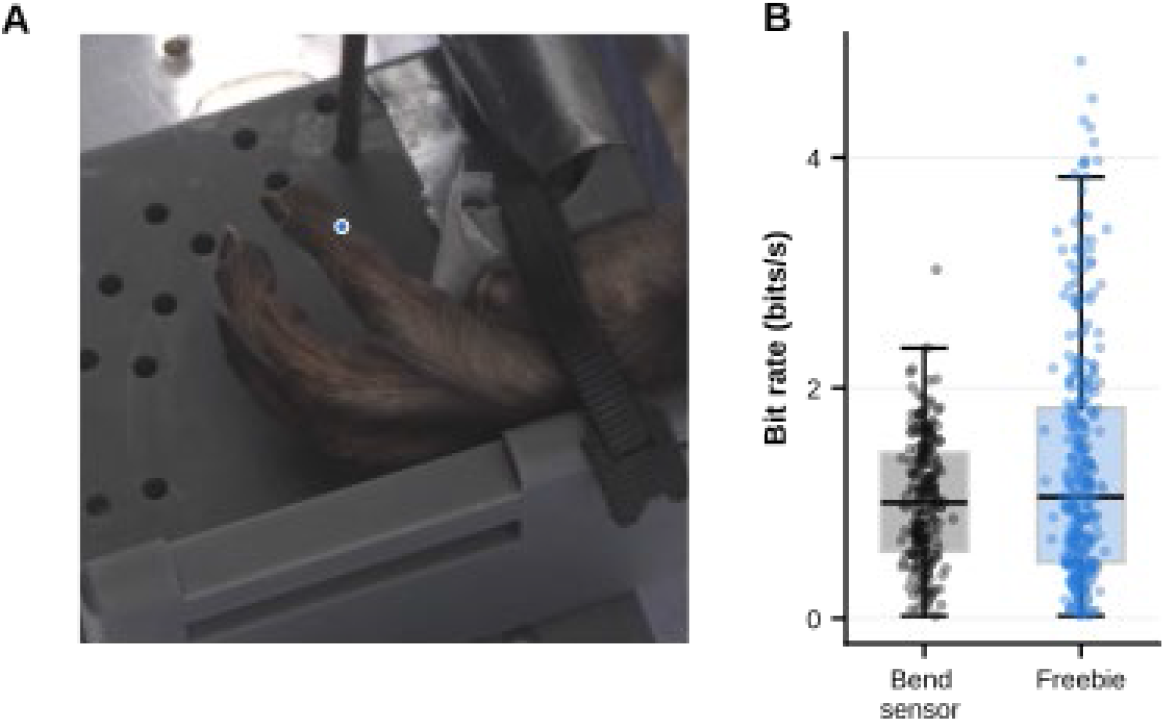
Freebie hand control and decoding without the manipulandum. (**A**) One camera frame from Freebie hand control with the live Freebie keypoint (index DIP) for that frame. Taken 4.2 seconds into the example trace in B, with the finger near full flexion (98%). (**B**) Bit rate of each successful trial. Groups: hand control with bend sensor (n = 232 trials, 1 run) and with Freebie (n = 320 trials, 2 runs); online KFs trained on bend-sensor data (n = 123 trials, 1 run) and on Freebie data (n = 123 trials, 1 run). Boxes show the median and interquartile range.

